# Linking host personality and parasitic infection: a meta-analysis

**DOI:** 10.1101/2025.06.30.662423

**Authors:** Anna Pia Piscitelli, Simone Messina, Lucas A. Wauters, Francesca Santicchia, Erik Matthysen, Herwig Leirs, Bram Vanden Broecke

## Abstract

Animal personality and parasite infections are key forces shaping the ecology and evolution of natural populations. Personality traits—such as activity, exploration, and boldness—shape how individuals interact with their environment and conspecifics, influencing both their exposure and susceptibility to parasite infection. In turn, parasites can impact host fitness and energy allocation, and may modify host behaviour either through manipulations to enhance transmission or as consequences of energetic trade-offs associated with mounting an immune response.

Despite growing interest in the interplay between behaviour and infection, the overall directionality and consistency of personality–parasite relationships remain unclear. This relationship is further modulated by ecological and biological factors, such as parasite type (e.g. micro-, ecto-, or endoparasites) and host type (e.g. intermediate versus definitive), which can influence both infection risk and the nature of behavioural responses. To disentangle these effects, we performed a meta-analysis of 226 effect sizes across 80 studies, assessing (i) the impact of experimental infections on host personality traits, and (ii) the correlation between personality and infection status in observational studies of wild populations—while accounting for variation in parasite groups and host roles.

In experimental studies, infected hosts exhibited significantly reduced levels of activity and exploration, while effects on boldness and aggressiveness were non-significant. These findings suggest that infection imposes energetic costs that suppress behaviours requiring sustained effort, such as movement and exploration. Conversely, observational studies showed a positive association between activity–exploration and infection probability, likely reflecting greater exposure of more active individuals to parasites via increased interaction with conspecifics or contaminated environments.

Meta-regression analyses further revealed that parasite type and host role modulate personality–infection dynamics. In experimental studies, microparasites were associated with reduced boldness and activity-exploration, while endoparasites led to reduced activity– exploration—particularly in intermediate hosts. Notably, hosts showed significant behavioural suppression in experimental contexts, but not in observational studies, potentially indicating that behaviourally tolerant individuals are favoured in natural environments where personality traits relate directly to fitness.

Together, these findings underscore the importance of ecological context and study design in interpreting personality–parasite associations. Experimental infections tend to reveal the physiological costs of infection, while observational studies highlight behavioural traits that modulate infection risk. By integrating data across host types, parasite groups, and methodological approaches, our meta-analysis provides a more comprehensive understanding of how personality and infection interact. These insights contribute to a broader effort to link behavioural ecology with disease ecology, clarifying how individual variation in behaviour shapes—and is shaped by—host–parasite dynamics.

## I. INTRODUCTION

Parasite infections and animal personality both play crucial roles in shaping the dynamics of natural populations. Parasitism acts as a strong ecological force that can profoundly alter population structure, density, and stability by reducing individual fitness through effects on host survival, reproductive success, and overall energy allocation (Barber and Dingemanse, 2010; Jacques-Hamilton et al., 2017; Khokhlova et al., 2002; Lachish et al., 2011; Pigeault et al., 2018). Animal personality, defined by Réale et al. (2007) as consistent behavioural differences among individuals of the same population or species through time and contexts, has been found to influence individual fitness (Biro and Stamps, 2008; Boyer et al., 2010; Haave-Audet et al., 2022; Jacques-Hamilton et al., 2017; Réale et al., 2010).

Animal personality is also tightly linked to individuals’ movement behaviour, sociality and immunocompetence, which in turn affect the risk of encountering parasites and becoming infected (Sih et al., 2018). In this way, personality contributes to the heterogeneity in exposure, susceptibility, and consequently the transmission of parasites within and among populations (Barber et al., 2017, 2000; Barber and Dingemanse, 2010; Kortet et al., 2010; Vanden Broecke et al., 2021, 2019). At the same time, parasites can exert differential selective pressures on hosts mediated by the link between personality and risk of infection (Barber et al., 2017; Barber and Dingemanse, 2010; Klein et al., 2004; Santicchia et al., 2020). These bidirectional interactions suggest a complex feedback loop between behaviour and disease. Understanding these interactions is crucial for identifying the mechanisms that influence parasite exposure, infection risk, and transmission patterns in natural populations.

It is commonly thought that proactive individuals (e.g. more explorative, active, bolder, aggressive and/or more sociable) have a higher probability of encountering parasites and, consequently, become infected (Barber and Dingemanse, 2010; Bohn et al., 2017; Dizney and Dearing, 2013; Ezenwa et al., 2016). However, this is not always the case. For example, less active and bolder pumpkinseed sunfish (*Lepomis gibbosus*) showed a higher likelihood of infection by trematodes and cestodes, suggesting that different personality traits can independently increase the risk of infection by a given parasite (Gradito et al., 2024). Similarly, less explorative multimammate mice (*Mastomys natalensis*) were more infected with viruses (Vanden Broecke et al., 2019), helminths (Vanden Broecke et al., 2021) and bacteria (Vanden Broecke et al., 2023) than more explorative conspecifics. These contrasting findings reveal that proactive behaviour does not uniformly predict infection risk, pointing instead to a more complex and context-dependent relationship.

One of the factors that may underlie this variation among studies is the type of parasite involved as the mode of transmission determines which personality traits are more strongly associated with infection risk (Fig. 1).

**Figure 1.**
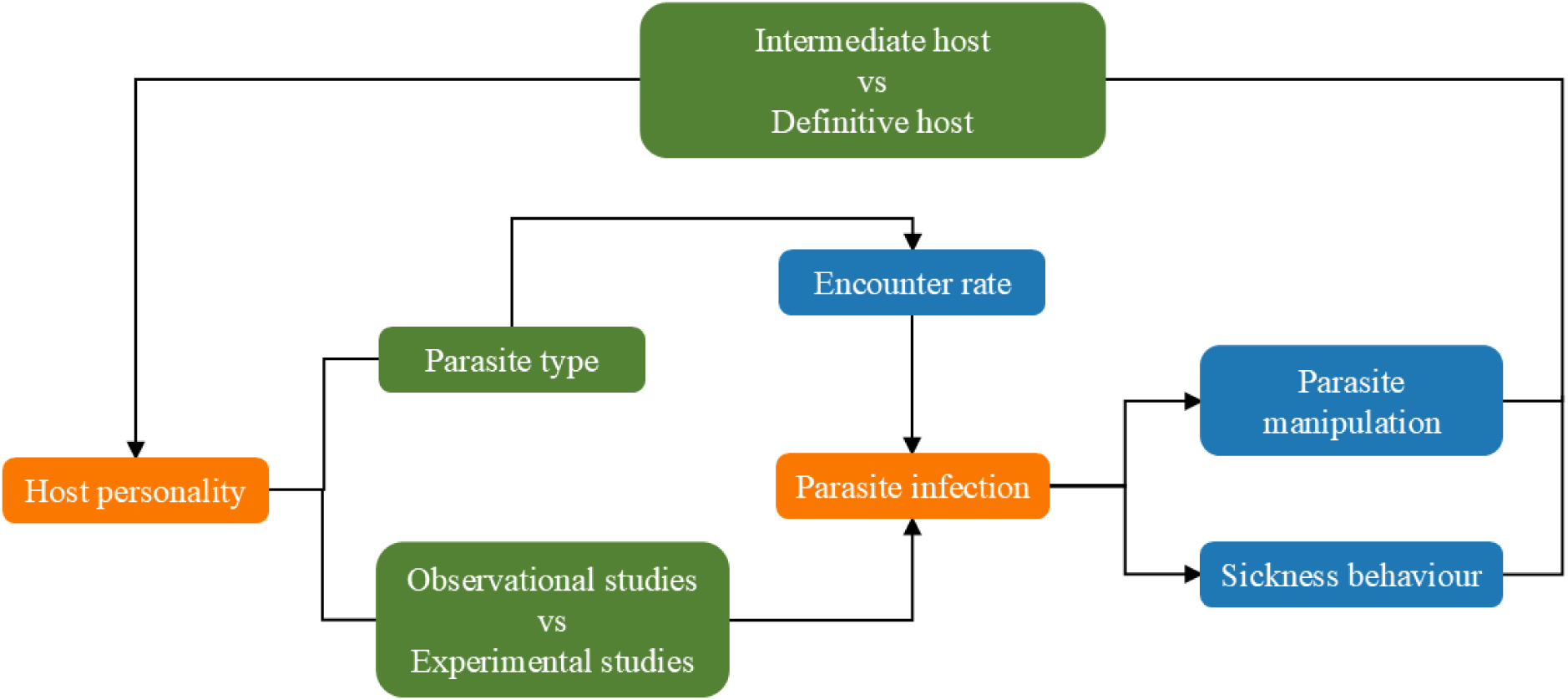
Schematic diagram showing the key conceptual relationships that connect individual personality differences with parasite infection. Orange boxes highlight the central research themes— personality and parasitic infection. Blue boxes represent the biological processes mediating the relationship between them, while green boxes indicate the key factors we propose as primary sources of heterogeneity across systems and studies.

Sociality and aggressiveness are expected to influence the infection risk of directly transmitted parasites, such as microparasites (e.g., viruses and bacteria), where contact rates with conspecifics are the primary transmission route (Klein et al., 2004; Petkova et al., 2018; Reisinger et al., 2015). In contrast, bolder and more explorative individuals are expected to have higher exposure risk to ectoparasites (e.g., fleas, ticks, mites) and endoparasites (e.g., helminths) that are transmitted indirectly through the environment (Patterson and Schulte-Hostedde, 2011; Rollins et al., 2021). This is because boldness and exploration are generally associated with increased mobility and a greater tendency to investigate novel environments, which raises the likelihood of encountering contaminated surfaces (Bohn et al., 2017; Boyer et al., 2010; Gyuris et al., 2016; Santicchia et al., 2019). Yet, to the best of our knowledge, no studies have systematically evaluated whether the link between animal personality and infection risk varies according to the type of parasite or the specific personality traits involved.

Moreover, the relationship between parasitic infection heterogeneity and personality is not solely driven by the effect of personality on encounter rates with parasites. Experimental studies have shown that parasites can actively manipulate host behaviour, leading to personality changes that enhance the parasite’s survival, reproduction, and transmission (Barber and Dingemanse, 2010; Kortet et al., 2010; Poulin, 2013). This is particularly evident for trophically transmitted parasites such as *Toxoplasma gondii* which alters mouse behaviour by reducing overall activity and neophobia, thereby increasing the likelihood that infected rodents are preyed by cats, which are the parasite’s definitive hosts (Piekarski, 1981; Webster, 2007).

The extent to which the hosts’ behaviour is affected by parasitic infection may vary depending on the role the host plays in the parasite’s life cycle. Intermediate hosts are more frequently subjected to specific behavioural modifications that enhance parasite transmission to the final host. For instance, a parasite may alter risk-taking or exploratory behaviour—either increasing or reducing it—in its intermediate host to enhance vulnerability to predation by the definitive host (Reisinger et al., 2015; Webster, 2007). Additionally, when the intermediate host is not yet infectious, the parasite may also reduce the host’s overall activity to lower predation risk and delay transmission until the appropriate stage (de Vries and van Langevelde, 2018; Dianne et al., 2011). Differently, behavioural changes in definitive hosts tend to be subtler, promoting parasite survival and reproduction rather than transmission (Benesh et al., 2021). Recognizing these distinctions is essential for accurately predicting infection outcomes and parasite spread in natural systems.

Nevertheless, not all behavioural changes in infected hosts result from active parasite manipulation to enhance its survival and transmission. From a physiological perspective, parasite infections can change the energy status of the host inducing a sickness effect with consequences for the expression of personality traits —a phenomenon known as “sickness behaviour” (Adelman and Martin, 2009; Kortet et al., 2010). Host resistance to parasite infections (i.e. the ability to limit within host pathogen replication) relies on innate and adaptive immune responses, which involves energy-demanding processes (Allen and Maizels, 2011; Grencis et al., 2014; Kohlmeier and Woodland, 2009; Medzhitov et al., 2012). These immune costs are associated with a diversion of nutritional energy away from other processes. According to the pace-of-life hypothesis such energy reallocation could instigate a trade-off between host resistance and energetically costly behaviours (Careau et al., 2008; Douhard et al., 2025; Réale et al., 2010). Disentangling whether personality changes reflect adaptive manipulation or physiological constraints remains a major challenge in host–parasite research. Despite an increasing number of studies exploring the links between personality and parasitism, no comprehensive quantitative synthesis has yet been conducted to assess their overall patterns and moderators. Studies investigating this correlation in wild populations are valuable to assess how different personality traits are linked to the risk of infections, however, the direction of causality remains uncertain. Experimental studies, on the other hand, provide a clear understanding by directly manipulating infection status, allowing researchers to isolate the effects of parasites on host behaviour. However, results from experimental studies may not always translate well to wild populations due to the controlled environments in which they are conducted.

Accounting for parasite type, host role, and specific personality traits is essential to understanding the complex and context-dependent nature of personality–disease dynamics. While identifying broad patterns across systems is valuable, overlooking such sources of heterogeneity can be misleading and lead to oversimplified conclusions. A comprehensive synthesis of data from diverse ecological contexts and study designs is therefore crucial to distinguish consistent trends from context-specific outcomes driven by ecological or methodological differences.

We conducted the first meta-analysis to assess how animal personality traits correlate with parasitic infection across diverse taxa and parasite types. Building on evidence that personality can both influence encounter rates and be altered by parasites, we investigated (i) how personality traits—such as boldness, exploration, or aggressiveness— correlate with infection and (ii) how they are affected by induced infections. To achieve this, we built two different datasets, one based on observational studies and one based on experimental studies. Furthermore, we examined whether variation in parasite type (micro-, ecto-, or endoparasites) and host type (intermediate or definitive) helps explain the observed heterogeneity in the relationship between personality traits and parasitic infections. By testing these moderating variables, our study provides novel insight into the ecological interplay between host behaviour and infection dynamics.

## II. MATERIALS AND METHODS

### 1) Literature screening

We conducted a systematic literature review of experimental and correlative studies linking animal personality and parasitic infections. Paper selection was conducted according to the standards outlined in the Preferred Reporting Items for Systematic reviews and Meta-Analysis (PRISMA; Rethlefsen et al. 2021; Figure S1). The literature screening was performed in Web of Science and Scopus, including every year up to November 2024, when the last search was conducted. We used a combination of behaviour and parasite-related keywords: “behavio$r*”, “personalit*”, “temperament*”, “behavio$ral syndrome*”, “socia*”, “explor*”, “activ*”, “bold*”, “aggress*”, “avoid*”, “shy*”, “parasit*”, “pathogen*”, “disease*”, “infect*”, “infest*”. We also excluded from the research human-related terms and other recurring terms unrelated to the focus of our study using NOT followed by the terms: “brood paras*”, “parasitoid*”, “feeding behavio$r*”, “mating behavio$r”, “medicine”, “HIV”, “AIDS”, “COVID*”, “diabet*”, “patient*”, “clinic*”, “cancer”, “Alzheimer”, “dementia”, “psycho*”, “Parkinson*”, “drug*”, “depression”, “schizophrenia”, “wom$n”, “m$n”, “adolescent*”. After initial searches, we added specific exclusion categories present in Web of Science and Scopus to avoid unrelated categories overwhelming the search results (see the Supplementary material).

After combining the results from Web of Science and Scopus and deleting duplicates (n= 675), the search yielded 7018 articles. Of these, 6773 articles were excluded at the title/abstract screening stage because they were: (i) human related studies, (ii) reviews, (iii) immune challenge experiments, (iv) studies on behaviours not related to our study (e.g. feeding behaviour, dominance, social network). We then carried out a full-text screening of the remaining 245 articles checking if they respected our inclusion criteria.

### 2) Inclusion criteria

We selected the studies based on the following criteria:

a) The article focused on at least one of the five categories of behaviours described by Réale et al. (2007) (i.e. activity, exploration, boldness, aggressiveness and sociability), including vertebrate and invertebrate species.
b) The article focused on animal personality, measured as a continuous variable to enable the calculation of correlation coefficients.
c) Personality is by definition repeatable and we initially aimed to include only studies that actually reported repeatability, however, very few of the studies reported it, nor cited previous work that documented repeatability in the same population. Therefore, to ensure an adequate sample size, we adopted a broader inclusion. We included articles where the behaviours were interpreted by the authors as personality trait or where the behaviours analysed were attributable to the personality traits defined by Réale et al. (2007), and where behaviours were measured using common tests for personality (e.g. open field test, hole board test, Y-maze, arena test, latency of emerging or resuming activities, social challenge and mirror image stimulation).
d) The article mentioned the parasite species or at least the type of parasite (e.g. ectoparasites, helminths, virus ect.) and the related measurement (i.e. presence and/or intensity of infection). When an article reported both measurements, they were counted as two distinct measurements (i.e. effect sizes).
e) The study reported sample size and statistical information so that the effect sizes could be calculated: mean and standard deviation or standard error, F from one-way anova, t-statistic, Z-statistic, chi-squared value, Pearson correlation coefficient, Point biserial correlation coefficient, Spearman correlation coefficient or standardized β-coefficient. When samples sizes, means and standard deviations or standard error were not available in the text, we extracted these information from the graphs using the software GetData Graph Digitizer (Fedorov et al., 2014). When basic statistics like sample size, means or standard errors were missing, authors were contacted and, if they didn’t respond, the articles were excluded.

Following these criteria, a total of 80 articles were retained.

### 3) Data coding

Articles were divided in two datasets. 1) The Experimental dataset included all the studies in which animals (captive or wild) were experimentally infected and/or infections were experimentally reduced or prevented through antiparasitic treatment, to assess the effect of infection on their behaviour. In these studies, the behaviour was compared between a control and treatment groups or within the same group before and after the infection/treatment. 2) The Observational dataset included studies investigating the link between infection status and behaviour, in free-living populations where individuals were naturally infected prior to behavioural assay. For each study, we noted the year of publication, host and parasite species name, type of parasite and type of host.

Regarding the personality traits, given the variation in terminology across studies, we applied standardized definitions of personality traits to ensure consistent data extraction and comparability, following the definitions made by Garamszegi et al. (2013) and Réale et al. (2007). We classified the behaviours in: I) **Boldness**, an individual’s reaction to potentially threatening situations (e.g., predator exposure), including measures like latency to resume normal activity or proximity to a threat; II) **Activity-Exploration**, any variable that measures the general level of activity of an individual (e.g. distance moved, time spent moving), including new situations; III) **Aggressiveness**, antagonistic behaviours directed toward conspecifics or mirror images, excluding dominance; IV) **Sociability**, any variable that measures an individual’s reaction to a conspecific (excluding aggression, aggregation, shoaling behaviour and social networks).

Parasites were classified in microparasites (e.g. viruses, fungi, bacteria, protozoan), ectoparasites (e.g. fleas, mites, ticks), and endoparasites (e.g. helminths). Hosts were classified as intermediate when the parasite was in a non-reproductive or asexual developmental stage, and as definitive when the parasite reached its adult, sexually mature stage and typically reproduced. This second classification also included endoparasites with environmental or multi-host transmission cycles.

### 4) Effect size calculation

In order to quantify the relationship between personality and parasite infection we used the Pearson’s correlation coefficient (r) as the measure of effect size. The correlation coefficient was calculated using the formulas provided in Koricheva et al. (2013) enabling comparison across studies. When standardized β-coefficient was available, it was directly interpreted as Pearson correlation coefficient r (Nieminen 2022). We then applied Fisher’s z-transformation to normalize the distribution of the response variable calculating *Zr* = 1/2 *ln*((1 + *r*) ∕ (1 − *r*)) and its variance *VZr* = 1 ∕ (*n* − 3) (Koricheva et al., 2013). Since r can range from -1 to +1 we assigned the direction of the effect size based on the association between behavioural traits and parasitic infection. Positive effect sizes were assigned when individuals exhibiting higher levels of activity, exploration, boldness, sociability, or aggressiveness also had higher parasite prevalence or load compared to their less active, explorative, bold, sociable, or aggressive conspecifics. Conversely, negative effect sizes were assigned when individuals with higher behavioural trait scores had lower parasite prevalence or load.

### 5) Meta-analytic technique

We conducted meta-analytic, multilevel, mixed-effects modelling using the *rma.mv* function from the *metafor* package (Viechtbauer, 2010) in the R environment (R Core Team, 2024). In our models, we included *Study ID*, *Effect Size ID*, and *Phylogeny* as random effects. *Study ID* accounted for the non-independence of multiple effect sizes derived from the same study, while *Effect Size ID* controlled for within-study effect size variation beyond sampling error (i.e. presence of more than one effect size within the same study). *Phylogeny* was incorporated to account for the shared evolutionary history between species (Cinar et al., 2022). To construct the phylogenies, we utilized the *rotl* package (Michonneau et al., 2016), which retrieves phylogenetic relationships from the Open Tree of Life. Branch lengths of phylogenetic trees were estimated using the *compute.brlen* function from the *ape* package (Paradis & Schliep 2018), and phylogenetic relatedness was modelled as a variance-covariance matrix. Initially, we also included *Species* and *Transmission* as random effects, but they did not improve the model’s AIC, so we excluded them and report the best-fitting models.

In our database some studies included effect sizes based on shared control groups (i.e., studies with one control group and multiple infection treatments). To address this non-independence, we weighted the effect sizes based on a variance-covariance matrix (Messina et al., 2023; Noble et al., 2017) and we used it instead of *VZr* in the models where shared control groups were present. We also calculated overall heterogeneity (I²Total) and the proportion attributed to each random factor using the i2_ml function (Nakagawa et al., 2020). I² reflects the percentage of variance among effect sizes not explained by sampling variance, with values above 75% indicating high heterogeneity, 50% as medium, and 20% as low (Higgins et al., 2003).

### 6) Meta-regression models

To assess the directionality of the relationship between personality and parasites, i.e. to determine whether personality influenced parasite infection or whether it was the infection that influenced personality, we ran a model testing for the overall effects of animal personality on infection using experimental and observational datasets separately. We included animal personality as a single moderator.

To assess whether parasite group (ectoparasite, microparasite, endoparasite) influences the relationship between animal personality and parasite infection, we ran separate models for each parasite group. We decided to subset the dataset because incorporating interactions in the model would have produced interactions with a limited number of effect sizes for reliable analysis. We used animal personality as moderator in independent models for each subset of interest (i.e. ectoparasite, microparasite, endoparasite). We set the minimum number of studies and effect sizes to include in each model at 3 for each personality type, including both papers measuring presence and intensity of infection. For this reason, we could not include all the personality traits in all models.

Finally, to assess whether infection has a differential impact on personality between intermediate and definitive hosts, we ran a model including host type as a moderator in both experimental and observational datasets. We focused specifically on endoparasites and on Activity-Exploration due to the limited number of studies available on the other personality traits.

### 7) Publication bias

We tested for potential publication bias by performing funnel plot analysis on the whole database. To this end, we plotted effect sizes on the y-axis and the inverse of the sample standard error, that is a measure of precision, on the x-axis (Nakagawa et al., 2022). We calculated precision from the variance of *Zr*. Then we tested the significance of the asymmetry using a multilevel version of Egger’s regression (Nakagawa et al., 2022). We also included as fixed effect *VZr* (the sampling variance of *Zr*) and as random effect *Study ID*, *Effect Size ID* and *Phylogeny*. Finally, we assessed the presence of a time lag effect in the publication of results by regressing standardized effect sizes (Zr) against publication year (Nakagawa et al., 2022), including the same random effects as the Egger regression model.

## III. RESULTS

### 1) Descriptive information

Overall, our work included 226 effect sizes from 80 studies and 67 species (Table S1). Vertebrates were the most represented taxon with 42 species and 168 effect sizes while invertebrates included 25 species and 58 effect sizes. Among vertebrates, mammals were the most represented (12 species and 89 effect sizes), followed by fish (13 species and 45 effect sizes), birds (6 species and 20 effect sizes), amphibians (8 species and 11 effect sizes), and reptiles (3 species and 3 effect sizes). Among invertebrates, Malacostraca (lobsters, shrimps and amphipods) were the most represented taxon (13 species and 33 effect sizes), followed by insects (7 species and 15 effect sizes), gastropods (2 species and 5 effect sizes), clitellates (1 species and 3 effect sizes), copepods (1 species and 1 effect size) and arachnids (1 species and 1 effect size). The dataset on observational studies included 40 studies, 132 effect sizes and 38 species (23 vertebrates and 15 invertebrates), while the dataset on experimental studies included 40 studies, 94 effect sizes and 33 species (21 vertebrates and 12 invertebrates).

Endoparasites were the most studied parasite group in both observational and experimental studies (46 articles: 24 observational and 22 experimental) followed by microparasites (24 articles: 12 observational and 12 experimental) and ectoparasites (11 articles: 5 observational and 6 experimental).

### 2) Meta-regression models

Models testing the overall relationship between infections and animal personality showed contrasting results between the experimental and observational datasets, in particular for Activity-Exploration. Results based on the experimental dataset showed significantly reduced active and explorative behaviours in infected individuals (estimate ± SE= -0.25 ± 0.13, CI= -0.50, 0.00) (Fig. 2A). In contrast, results based on the observational dataset showed a positive correlation between Activity-Exploration and infection (estimate ± SE= 0.13 ± 0.06, CI= 0.00, 0.25) (Fig. 2B). Similar opposite patterns between databases were observed for Boldness and Aggressiveness, although none of the associations were found to be significant (Fig. 2A-B).

**Figure 2.**
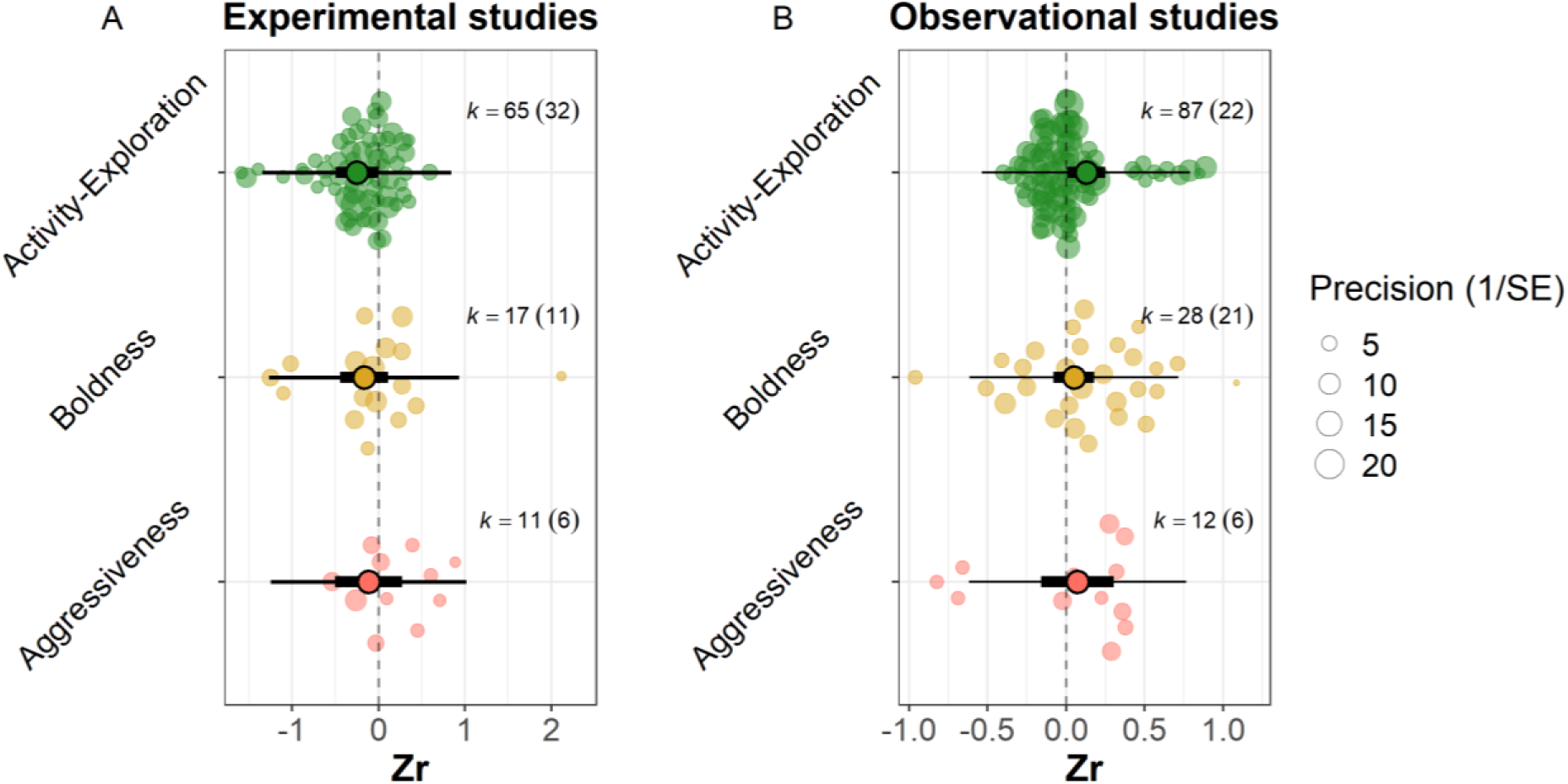
Mean estimates (Zr) of the relationship between individual personality and parasite infection are presented for both the A) Experimental and B) Observational studies. Opaque circles represent the effect sizes, with their size corresponding to precision (1/SE). Bold lines indicate 95% confidence intervals, while thin lines represent 95% prediction intervals. The symbol ’k’ denotes the number of effect sizes, with the number of studies indicated in parentheses.

The models testing for effects of parasite types on personality showed reduced Activity-Exploration in hosts experimentally infected with endoparasites (estimate ± SE= -0.18 ± 0.08, CI= -0.33, -0.02) (Fig. 3A), while Boldness showed no significant association (Fig. 3A). A significant reduction in Boldness was found in hosts experimentally infected with microparasites (estimate ± SE= -0.86±0.36, CI= -1.63, -0.09) (Fig. 3C), whereas Activity-Exploration showed no significant correlation with ectoparasites (Fig. 3E).

**Figure 3.**
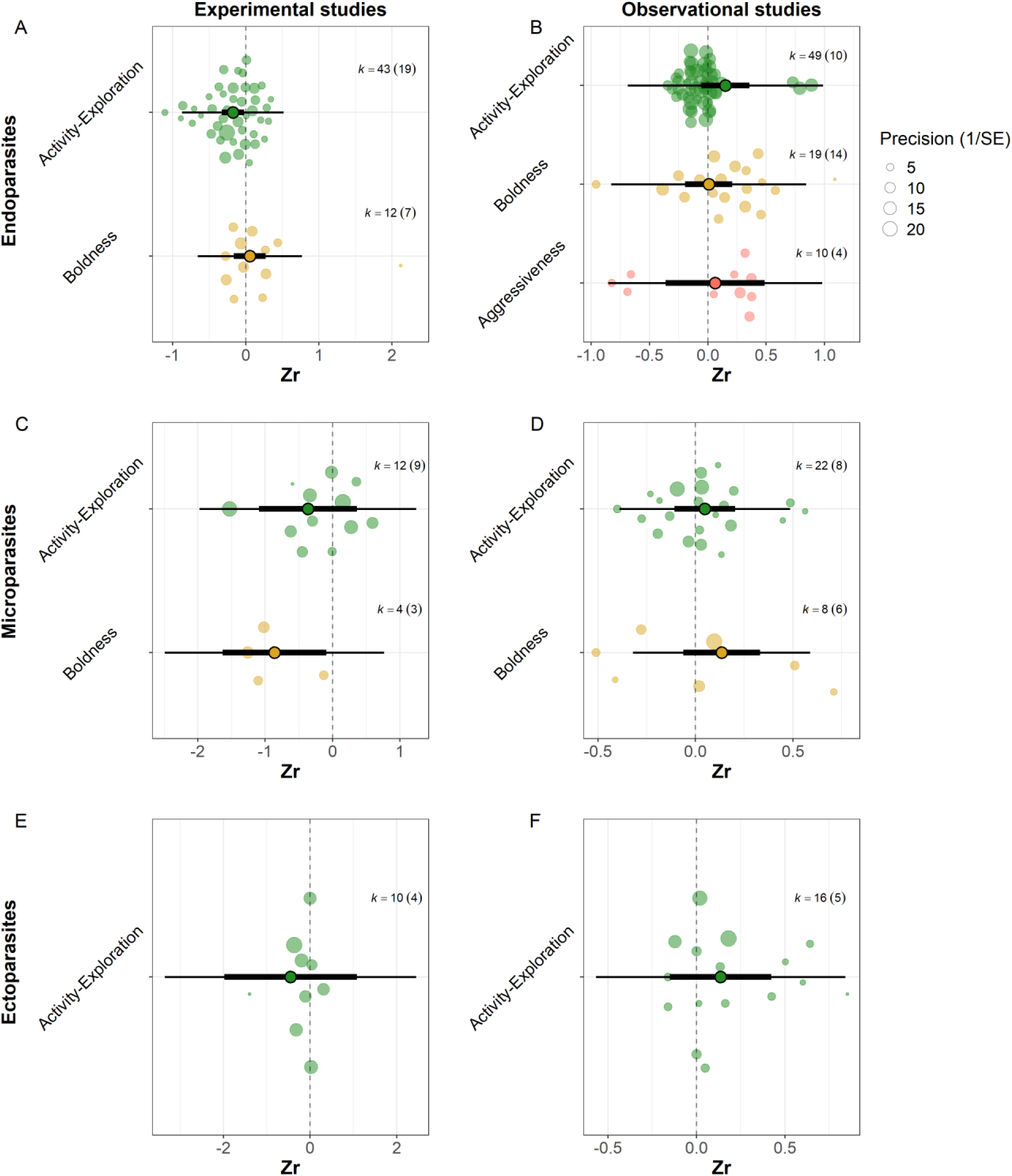
Mean estimates (Zr) of the relationship between individual personality and A-B) endoparasite, C-D) microparasite and E-F) ectoparasite infection are presented for both the Experimental and Observational studies. Opaque circles represent the effect sizes, with their size corresponding to precision (1/SE). Bold lines indicate 95% confidence intervals, while thin lines represent 95% prediction intervals. The symbol ’k’ denotes the number of effect sizes, with the number of studies indicated in parentheses.

In observational studies, we found a marginally positive correlation between Activity-Exploration and endoparasite infection (estimate ± SE= 0.15±0.10, CI= -0.05, 0.35) (Fig. 3B) while it was not significant for micro- and ectoparasites. Aggressiveness and Boldness did not exhibit significant patterns in any of the models across endo-, micro- and ectoparasites (Fig. 3B-D-F).

Finally, for intermediate hosts, we found a significant negative correlation between Activity-Exploration and endoparasite infection in experimental studies (estimate ± SE= - 0.25±0.09, CI= -0.44, -0.06) (Fig. 4). In observational studies, the relationship between host type and parasite infection was not significant for either intermediate or definitive hosts (Fig. 4).

**Figure 4.**
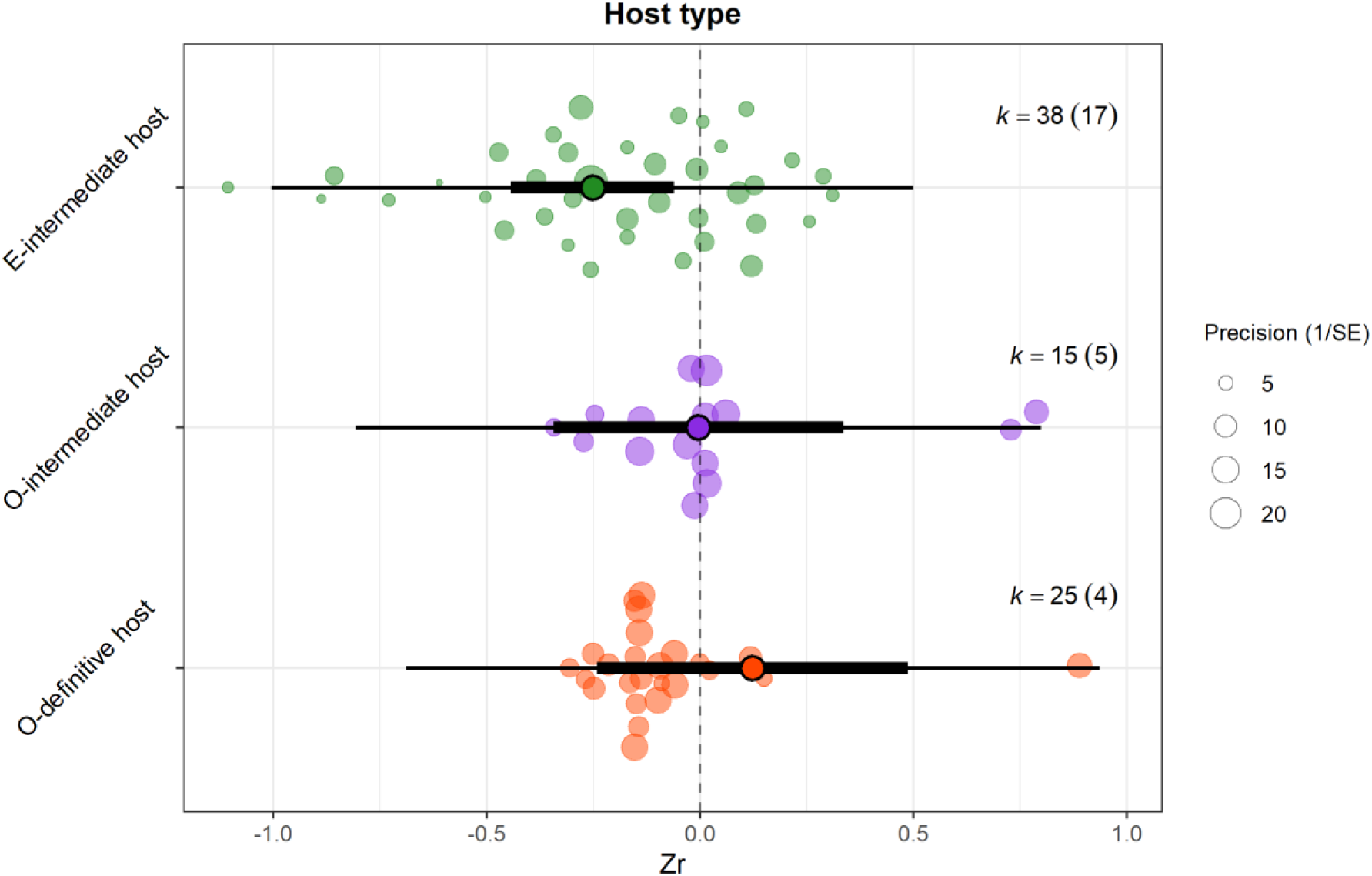
Mean estimates (Zr) of the relationship between Activity-Exploration in intermediate and definitive hosts and endoparasites infection. Green circles represent intermediate hosts in experimental studies (E), while violet and red circles denote intermediate and definitive hosts in observational studies (O), respectively. Opaque circles represent the effect sizes, with their size corresponding to precision (1/SE). Bold lines indicate 95% confidence intervals, while thin lines represent 95% prediction intervals. The symbol ’k’ denotes the number of effect sizes, with the number of studies indicated in parentheses.

### 3) Publication bias

We found no evidence of publication bias from funnel plot visualization (Fig. S2). Similarly, Egger’s regression test indicated no publication bias, as the intercept was not significantly different from zero (estimate ± SE= -0.05±0.08, p = 0.51, CI=-0.22, 0.11) and the coefficient for standard error was not significantly different from zero (estimate ± SE= 0.02±0.39, p = 0.96, CI= -0.74, 0.78). Additionally, we detected no time-lag bias, as there was no significant relationship between effect size and publication year. The intercept of effect size was not statistically significant (estimate ± SE=11.10±12.82, p = 0.40, CI= -14.98, 37.20) and neither was the coefficient for the publication year (estimate ± SE= -0.01±0.01, p = 0.40, CI= -0.02, 0.01).

## IV. DISCUSSION

Free-living animals are frequently exposed to multiple parasites, and although animal personality has been implicated in host-parasite interactions, the direction and the moderators of this relationship remain unresolved. Through a meta-analytic approach, we showed that in observational studies more active and explorative individuals are more likely to be infected, in particular by endoparasites, indicating that such personality trait increases the risk of encountering such parasites in the environment. Furthermore, we found downregulated proactive behaviours (activity, exploration and boldness) following experimental infections, but depending on the type of parasite. These results suggest different trade-offs between immune response and energetically expensive behaviours, driven by the type of infection.

### 1) Experimental studies

Our analyses showed a negative correlation between the personality trait “Activity-Exploration” and parasite infection in experimental studies, indicating that hosts generally became less active and explorative after experimental infection. One possible explanation is that this reduction reflects parasite-induced behavioural manipulation aimed at enhancing transmission. This interpretation is partially supported by our finding that reduced activity was especially evident in intermediate hosts infected by endoparasites—parasites often associated with trophic transmission strategies that benefit from increasing the host’s susceptibility to predation by definitive hosts. However, behavioural changes do not always reflect adaptive manipulation. Many parasites, particularly those transmitted through environmental exposure, may alter host behaviour as a non-adaptive consequence of infection. Since our dataset encompassed a broad range of parasite species, including those with environmental transmission, the overall reduction in activity and exploration likely reflects physiological costs of infection—such as immune activation and energy depletion—rather than active behavioural manipulation.

These effects may be further amplified in experimental studies, where high parasite loads are often employed and where behaviours are often measured shortly after infection to ensure measurable outcomes, intensifying the impact on host behaviour.

This downregulation of the behaviour was particularly evident in infections caused by microparasites and endoparasites, with boldness significantly decreasing and activity– exploration showing a declining trend following microparasite infections, while activity– exploration decreased in response to endoparasite infections. Microparasites, such as bacteria, viruses, and protozoa, often induce acute infections characterized by rapid onset and heightened immune response which is metabolically demanding (Kohlmeier and Woodland, 2009; Martin et al., 2003). Therefore, reduced boldness and activity-exploration may result from systemic sickness behaviours, such as withdrawal and reduced responsiveness, that limit risky or energetically costly behaviours during infection.

Similarly, endoparasites, which reside within the host’s body, often place a significant energetic burden on their hosts by consuming resources, disrupting physiological processes, or triggering immune responses (Brosschot and Reynolds, 2018; Grencis et al., 2014; Shanebeck et al., 2022). However, many endoparasites actively modulate the host’s immune system— often downregulating both humoral and cell-mediated responses to promote their persistence (Grencis et al., 2014). As a result, the behavioural effects of endoparasites may stem more from chronic nutritional drain than from immune-mediated sickness behaviours. This energetic cost is likely to have a greater impact on continuous traits like activity and exploration, which require sustained energy expenditure. In contrast, boldness—characterized by short-term responses to risk or novelty—may be less affected by resource depletion and more sensitive to immuno-physiological disruption.

In contrast, the effect on host behaviour appears less pronounced for ectoparasites. Ectoparasites often establish transient infections with relatively low physiological impact on the host which can also vary strongly among host and parasite species (Brophy and Luong, 2022; Careau et al., 2010; Heylen et al., 2010; Heylen and Matthysen, 2008; Rynkiewicz et al., 2013). For example, in white-footed mice (*Peromyscus leucopus*), higher bacterial killing ability was positively associated with tick burden but showed no correlation with flea infestation (Rynkiewicz et al., 2013). Nonetheless, the apparent lack of a consistent pattern in our analysis may also reflect insufficient data on ectoparasite infections, rather than the absence of physiological effects.

### 2) Observational studies

In observational studies, we found an increased infection rates in more active and explorative individuals. In the wild, more active and explorative individuals may have greater exposure to parasites due to increased movement and broader space use, which increase the likelihood of encountering sources of infection (Barber and Dingemanse, 2010; Boyer et al., 2010; Ezenwa et al., 2016). For instance, immigrant great tits (*Parus major*) show a stronger exploratory tendency than residents (Korsten et al., 2013; Quinn et al., 2011), and more explorative male red squirrels (*Sciurus vulgaris*) use larger core-areas within their home range (Wauters et al., 2021). This high exposure increases the probability of encountering parasites that may be present in soil, water, vegetation, or in environments contaminated by the excretions of infected animals (Bohn et al., 2017; Dunn et al., 2011; Marinov et al., 2017).

A positive correlation between Activity-Exploration and infections probability may occur for endoparasites and ectoparasites which can be transmitted through environments. Nonetheless, while we found a marginally positive correlation between active and explorative levels and endoparasite infections, no significant correlation was found with ectoparasites.

Ectoparasite’ infection success may not be solely affected by host behaviours but these parasites can actively select their hosts and rely on attachment and feeding strategies that are less dependent on the host’s behaviour (Christe et al., 2007; Fracasso et al., 2019). As a result, the influence of Activity-Exploration on ectoparasite interactions may be limited and/or highly dependent on the exposure rate and morphological (e.g. body size) or physiological conditions of the host species under study. Additionally, as mentioned above, ectoparasites themselves might have a limited impact on their host behaviour compared to other types of parasites.

In contrast, endoparasite infections in the wild—particularly those caused by helminths—are typically acquired through repeated, low-dose exposures over time (Grencis et al., 2014), with more active and exploratory individuals being at greater risk due to increased chances of encountering parasites in the environment. This repeated low-dose exposure often elicits a modulated immune response rather than a highly inflammatory one, enabling hosts to tolerate the parasite without experiencing severe pathology that might otherwise impair behaviour (Adelman and Hawley, 2017; Grencis et al., 2014; Medzhitov et al., 2012).

Notably, these results contrast with those from experimental studies, where infection was associated with reduced activity–exploration. This discrepancy may reflect the greater ecological relevance of personality traits under natural conditions. Behaviours such as activity, exploration, and boldness are tightly linked to individual fitness in the wild, as they are associated with dominance status, increased access to resources and early reproduction (Biro and Stamps, 2008; Snijders et al., 2017). In wild-conditions, reducing such behaviours in response to infection may impose high fitness costs. Therefore, in this context, wild animals may exhibit behavioural tolerance—the maintenance of normal behavioural patterns despite infection—as a strategy to mitigate these costs associated with an increase infection risk (Adelman and Hawley, 2017). Unlike resistance (which reduces parasite burden) or physiological tolerance (which minimizes damage caused by parasites), behavioural tolerance allows hosts to maintain fitness-enhancing behaviours while infected (Adelman and Hawley, 2017; Medzhitov et al., 2012). In contrast, captive animals, lacking natural selective pressures and ecological challenges, may be more prone to showing passive sickness behaviours, thereby amplifying the apparent impact of infection on personality traits.

### 3) Sociability and Aggressiveness

Our analysis revealed no significant correlation between aggressiveness and parasite infection, a pattern consistent across both experimental and observational studies. Unlike traits such as activity or exploration, which are expressed consistently across time and contexts, aggressiveness tends to occur in discrete, short-lived social interactions, often triggered by specific environmental or social stimuli (Réale et al., 2007). As a result, its overall energetic or immunological cost to the host may be lower or more context-dependent. This limited temporal expression reduces the likelihood that aggressiveness directly influences exposure to parasites or the physiological consequences of infection.

Sociability was not included in our meta-analysis due to the limited number of studies using standardised, repeatable behavioural assays. Most studies quantifying sociability relied on social network analysis (SNA), which we excluded for two main reasons. First, our inclusion criteria focused on personality traits measured through repeatable individual-based tests, such as mirror image stimulation, to ensure consistency across taxa and study types. Second, the link between parasitism and social network metrics has already been the focus of a recent meta-analysis (Briard and Ezenwa, 2021), which comprehensively explored how infection risk varies with social connectivity, centrality, and group structure. They reported a positive association between social network metrics and parasite infection suggesting that individuals engaging in more social interactions or sharing space with more conspecifics, face a higher risk of infection, although the high heterogeneity in effect sizes indicates substantial variation across studies.

## 4) Limitations and future directions

Overall, our findings suggest that, in natural systems, the direction of causality often flows from behaviour to infection, via increased exposure associated with traits such as activity and exploration. In contrast, the reverse pathway—from infection to behaviour through sickness effects or parasite manipulation—may be more subtle, transient, or masked by the capacity of the host to tolerate the infection. This highlights personality as a key driver of disease dynamics leading to a predictable variation in who become infected which is crucial for managing disease outbreaks in both wildlife and domestic settings. Moreover, behaviour-driven infection risk can affect broader ecological processes influencing population structure, community composition, and trophic interactions, particularly by increasing the vulnerability of more exposed individuals to predation or fitness reductions. Nonetheless, caution is warranted when interpreting studies that measure personality and infection simultaneously in the wild, as behavioural traits may already be altered by infection. Indeed, as suggested by experimental studies, endoparasites and microparasites can alter host personality. However, to enhance their ecological relevance, experimental studies should aim to better replicate natural infection dynamics, particularly in terms of infection dose and timing and investigate how behaviour influence infection risk in a controlled environment.

Our results also highlight the importance of accounting for both parasite and personality trait types. Most studies in our dataset examined only one parasite type and focused on one or two personality traits, even though wild animals are frequently co-infected with multiple parasites and display a broad range of personality dimensions. Because different parasites may interact with different aspects of host behaviour, certain traits may be linked to infection risk or behavioural alteration for one parasite, but not another. To capture the full complexity of these interactions, it is crucial to assess a wider array of personality traits and infection types wherever possible. Notably, traits such as activity and exploration were overrepresented, compared to boldness and sociability. Furthermore, the great majority of experimental studies focused on intermediate hosts, thus limiting our understanding on the effects of parasitic infections on their final hosts.

Future research should address these limitations by integrating experimental and observational approaches to capture both pre- and post-infection dynamics. Studies should expand their focus to include a broader range of personality measurements, host taxa, parasite types, and ecological contexts. Moreover, incorporating immune responses and co-infection dynamics into personality research could shed light on how infections influence—and are influenced by—individual behavioural differences. By adopting a more holistic approach, future work can build a deeper understanding of the complex and multifaceted interactions that drive host-parasite relationships in natural systems.

## V. CONCLUSIONS

1) Contrasting patterns emerged between experimental and observational studies. Experimentally infected hosts exhibited significant reductions in activity and exploration, likely reflecting energetic trade-offs associated with immune activation. In contrast, observational studies revealed a positive association between activity– exploration and infection risk, suggesting that more active and explorative individuals may experience higher exposure but they might also tolerate the infection.
2) Parasite type shaped the personality–infection relationship. In experimental studies, microparasites reduced both boldness and activity–exploration, whereas endoparasites predominantly downregulated activity–exploration. In observational studies, only endoparasite infections showed a marginally positive correlation with activity–exploration.
3) Host type influenced behavioural responses to infection. Stronger effects of parasite infection on personality traits were observed in intermediate in experimental studies.
4) Our findings highlight the importance of combining experimental and observational methods, expanding the focus to a broader range of personality traits, parasite groups and host types, and incorporating eco-immunology to better understand the heterogeneity in disease dynamic.

## ACKNOWLEDGEMENTS

This research was supported by the scholarship (1105322N) and project grant (G065720N) awarded by the Research Foundation – Flanders (FWO).

