## Supplementary data for "Linking host personality and parasitic infection: a meta-analysis"

*Corresponding author:

**Search strategy**

We ran a systematic literature search, according to the standards outlined in the Preferred Reporting Items for Systematic reviews and Meta-Analysis (PRISMA; Rethlefsen et al., 2021). The research was performed in November 2024 in Web of Science and Scopus.

***Web of Science***

The search was conducted in "Topic" using the following keywords: (behavio$r* OR personalit* OR temperament* OR "behavio$ral syndrome*") AND (socia* OR explor* OR activ* OR bold* OR aggress* OR avoid* OR shy*) AND (parasit* OR pathogen* OR disease* OR infect* OR infest*) NOT (brood paras* OR parasitoid*) NOT "feeding behavio$r*" NOT "mating behavio$r" NOT (medicine OR HIV OR AIDS OR COVID* OR diabet* OR patient* OR clinic* OR cancer OR Alzheimer OR dementia OR psycho* OR Parkinson* OR drug* OR depression OR schizophrenia) NOT (wom$n OR m$n OR adolescent*). The search lead to 40026 articles. To limit the amount of articles we refined the search including only articles in English and this lead to 33643 articles. Then, through the "Research areas", we included (environmental sciences ecology, zoology, behavioural sciences, parasitology, veterinary sciences, biodiversity and conservation) and excluded (neurosciences neurology, public environmental occupational health, life sciences biomedicine other topics, agriculture, biochemistry molecular biology, genetic heredity, engineering, toxicology, endocrinology metabolism, water resources, science technology other topics, pharmacology pharmacy) categories lead to 4799 articles.

***Scopus***

The search was conducted in "Article titles, abstract, keywords" using the following keyworkds: (behavio?r* OR personalit* OR temperament* OR "behavio?ral syndrome*") AND (socia* OR explor* OR activ* OR bold* OR aggress* OR avoid* OR shy*) AND (parasit* OR pathogen* OR disease* OR infect* OR infest*) AND NOT (brood paras* OR parasitoid*) AND NOT "feeding behavio?r*" NOT "mating behavio?r" AND NOT (medicine OR HIV OR AIDS OR COVID* OR diabet* OR patient* OR clinic* OR cancer OR Alzheimer OR dementia OR psycho* OR Parkinson* OR drug* OR depression OR schizophrenia) AND NOT (wom?n OR m?n OR adolescent*). The search resulted in

12144 articles. To limit the amount of articles we included only articles in English and we got 8783 papers. Then, using the ”Subject areas” available in Scopus we included (Agricultural and Biological Sciences, Environmental Science, Veterinary) and excluded (Biochemistry genetic and molecular biology, medicine, neuroscience, mathematics, social sciences, pharmacology toxicology and pharmaceutics) categories and we got at the end 2894 articles.

Citations from Scopus and Web of Science (n=7693) were uploaded in EndNote and duplicates (=675) were removed. 7018 articles were screened for title and abstract and 6773 were excluded. After these step 245 papers were screened for full text.

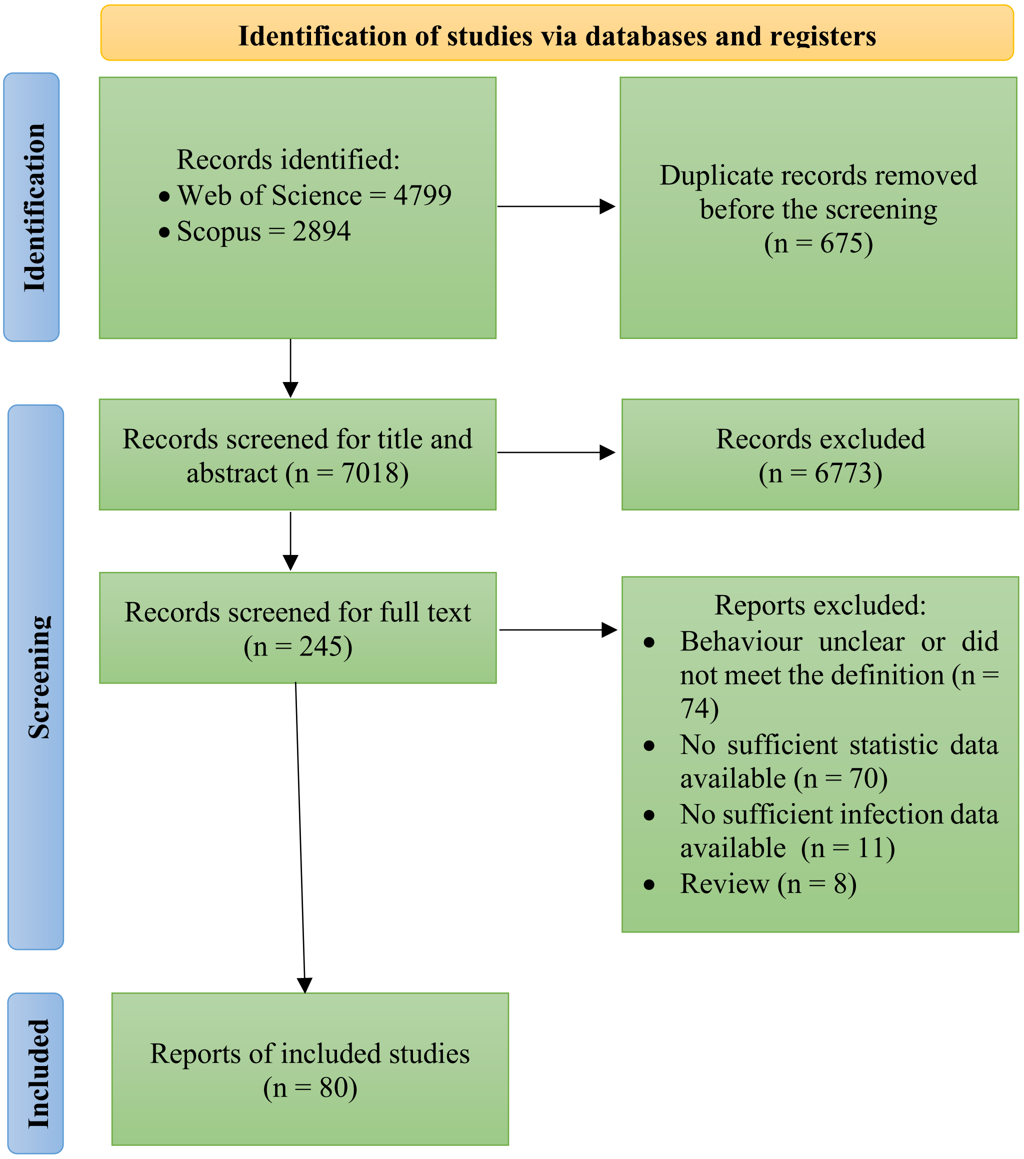

**Figure S1.** PRISMA flow-chart showing the number of articles retained at each phase of the systematic review and meta-analysis.

**Table S1.** List of species included in the meta-analysis, with relative taxonomic class, parasite and study citation.

| | Class | Species | Parasite | Study |
|---|---|---|---|---|
| **Vertebrata** | Actinopterygii | *Culaea inconstans* | endoparasite | McLennan & Shires 1995 |
| | Actinopterygii | *Etheostoma caeruleum* | ectoparasite | Crane et al. 2011 |
| | Actinopterygii | *Gasterosteus aculeatus* | microparasite | Petkova et al. 2018 |
| | Actinopterygii | *Neogobius melanostomus* | endoparasite | Flink et al. 2017 |
| | Actinopterygii | *Oncorhynchus mykiss irideus* | endoparasite | Gopko et al. 2015, 2017; Klemme et al. 2016; Mikheev et al. 2010 |
| | Actinopterygii | *Oryzias latipes* | microparasite | Vindas et al. 2023 |
| | Actinopterygii | *Phoxinus phoxinus* | endoparasite | Kortet et al. 2015 |
| | Actinopterygii | *Pimephales promelas* | endoparasite | Pan et al. 2016; Shirakashi & Goater 2002 |
| | Actinopterygii | *Poecilia vivipara* | endoparasite | Santos et al. 2011; Santos & Santos 2013 |
| | Actinopterygii | *Poeciliopsis retropinna* | endoparasite | Hagmayer et al. 2020 |
| | Actinopterygii | *Lepomis gibbosus* | endoparasite | Gradito et al. 2024 |
| | Actinopterygii | *Squalius cephalus* | ectoparasite | Horky et al. 2014 |
| | Amphibia | *Ambystoma tigrinum stebbinsi* | microparasite | Parris et al. 2004 |
| | Amphibia | *Anaxyrus americanus* | endoparasite | Koprivnikar & Urichuk 2017 |
| | Amphibia | *Cyclorana australis* | endoparasite | Nelson et al. 2015 |
| | Amphibia | *Limnodynastes convexiusculus* | endoparasite | Nelson et al. 2015 |
| | Amphibia | *Litoria nasuta* | endoparasite | Nelson et al. 2015 |
| | Amphibia | *Pseudacris regilla* | endoparasite | Goodman & Johnson 2011 |
| | Amphibia | *Rana aurora* | microparasite | Lefcort &Blaustein 1995 |
| | Amphibia | *Rhinella marina* | endoparasite | Nelson et al. 2015 |
| | Aves | *Ficedula albicollis* | microparasite | Garamszegi et al. 2015 |
| | Aves | *Himatione sanguinea* | microparasite | Yorinks & Atkinson 2000 |
| | Aves | *Luscinia megarhynchos* | microparasite | Marinov et al. 2017 |
| | Aves | *Motacilla flava* | microparasite | Marinov et al. 2017b |
| | Aves | *Parus major* | ectoparasite microparasite | Dunn et al. 2011 ; Rollins et al. 2021 |

| | | | | |
|---|---|---|---|---|
| | Aves | *Passer domesticus* | endoparasite | Garcia-Longoria et al. 2015 |
| | Mammalia | *Dicrostonyx richardsoni* | endoparasite | Quin et al. 1987 |
| | Mammalia | *Mastomys natalensis* | endoparasite microparasite | Vanden Broecke et al. 2018, 2021, 2023 |
| | Mammalia | *Mus musculus* | endoparasite microparasite | Hrdà et al. 2000; Kavaliers & Colwell 1995; Skallova et al. 2006; Cox et al. 1998; Cox & Holland 2001 |
| | Mammalia | *Myotis lucifugus lucifugus* | ectoparasite microparasite | Webber et al. 2015; Wilkox et al. 2014 |
| | Mammalia | *Oryctolagus cuniculus* | ectoparasite endoparasite | Arias-Hernandez 2019; Hallal-Calleros et al. 2013 |
| | Mammalia | *Peromyscus leucopus* | Endoparasite ectoparasite | Cramer & Cameron 2007; Caron-Lévesque & Careau 2023 |
| | Mammalia | *Peromyscus maniculatus* | microparasite | Dizney & Dearing 2013 |
| | Mammalia | *Rattus norvegicus* | endoparasite microparasite | Blecharz-Klin et al. 2022; Klein et al. 2004 |
| | Mammalia | *Sciurus carolinensis* | endoparasite | Santicchia et al. 2019 |
| | | *Sciurus vulgaris* | endoparasite | Santicchia et al. 2020 |
| | | *Tamias minimus* | ectoparasite | Bohn et al. 2017 |
| | Mammalia | *Tamias striatus* | endoparasite | Patterson & Schulte-Hostedde 2011 |
| | Reptilia | *Iberolacerta cyreni* | endoparasite | Horvath et al. 2016 |
| | Reptilia | *Sceloporus occidentalis* | ectoparasite | Lanser et al. 2021 |
| | Reptilia | *Tropidurus hispidus* | endoparasite | Maia-Carneiro et al. 2018 |
| **Invertebrata** | Arachnida | *Agelenopsis pennsylvanica* | microparasite | Parks et al. 2018 |
| | Clitellata | *Erpobdella octoculata* | endoparasite | Karvonen et al. 2017 |
| | Copepoda | *Macrocyclops albidus* | endoparasite | Benesh 2019 |
| | Gastropoda | *Biomphalaria glabrata* | endoparasite | Alberto-Silva et al. 2015; Boissier et al. 2003 |
| | Gastropoda | *Littorina littorea* | endoparasite | Seaman & Briffa 2015 |
| | Insecta | *Ameletus similior* | endoparasite | Benton et al. 1990 |
| | Insecta | *Anopheles darlingi* | microparasite | Bastos et al. 2024 |
| | Insecta | *Drosophila nigrospiracula* | ectoparasite | Brophy & Luong 2021 |
| | Insecta | *Formica rufa* | microparasite | Turner & Hughes 2018 |
| | Insecta | *Periplaneta australasiae* | endoparasite | Moore et al. 1994 |
| | Insecta | *Pyrrhocoris apterus* | ectoparasite | Gyuris et al. 2016 |
| | Insecta | *Temnothorax nylanderi* | endoparasite | Scharf et al. 2012 |
| | Malacostraca | *Austridotea annectens* | endoparasite | Friesen et al. 2017; Hansen & Poulin 2005 |
| | Malacostraca | *Dikerogammarus haemobaphes* | microparasite | Bojko et al. 2019 |

| | | | |
|---|---|---|---|
| Malacostraca | *Faxonius rusticus* | endoparasite | Reisinger et al. 2015 |
| Malacostraca | *Gammarus duebeni celticus* | microparasite | Fielding et al. 2005 |
| Malacostraca | *Gammarus pulex* | endoparasite | Baldauf et al. 2007; Dianne et al. 2014 ; Franceschi et al. 2007 |
| Malacostraca | *Gammarus roeselii* | endoparasite | Médoc et al. 2009 |
| Malacostraca | *Orconectes propinquus* | endoparasite | Reisinger et al. 2015 |
| Malacostraca | *Orconectes virilis* | endoparasite | Reisinger et al. 2015 |
| Malacostraca | *Pacifastacus leniusculus* | ectoparasite | James et al. 2015 |
| Malacostraca | *Palaemon pugio* | endoparasite | Kunz & Pung 2004 |
| Malacostraca | *Pallaseopsis quadrispinosa* | endoparasite | Benesh et al. 2008 |
| Malacostraca | *Paracalliope fluviatilis* | endoparasite | Friesen et al. 2017 |
| Malacostraca | *Paracorophium excavatum* | endoparasite | Friesen et al. 2017 |

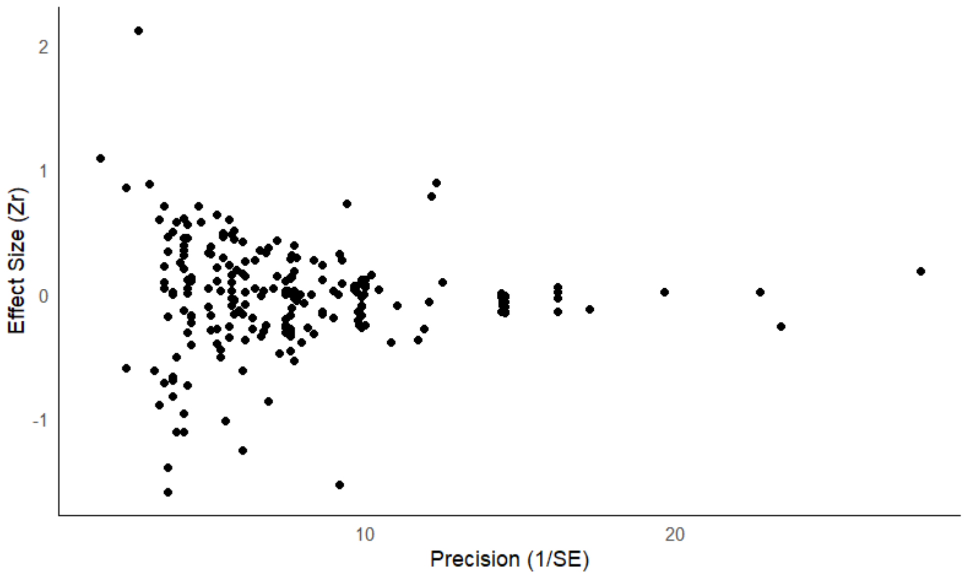


**Figure S2.** Funnel plot of effect sizes (y-axis) against the inverse of the sample standard error (x-axis). Data are referred to the full dataset including both experimental and observational studies.
